# Isolation and characterization of Bagaza virus: insights into host adaptability and zoonotic potential

**DOI:** 10.64898/2026.02.05.703981

**Authors:** Laura Albentosa-González, João Queirós, Catarina Fontoura-Gonçalves, David Gonçalves, Paulo Célio Alves, Armando Arias, Belén García-Navarro, Rosario Sabariegos, Ursula Höfle, Antonio Mas

## Abstract

Bagaza virus (BAGV) is an emerging orthoflavivirus that primarily infects wild birds, but its ability to infect other hosts remains poorly understood. We isolated BAGV RNA from the brain of a dead red-legged partridge and evaluated viral adaptation and replication kinetics in avian, mosquito, mammalian, and human cell lines. BAGV replicated efficiently in monkey, mosquito, duck, and selected human cell lines (HEK-293T and Huh7.5), whereas replication was limited in other human, hamster, and chicken fibroblast cell lines, indicating variable cellular susceptibility. These differences may reflect host-specific adaptation or distinct innate immune responses. Viral RNA replication complexes localized to the cytoplasm, as shown by immunocytochemical detection of dsRNA. Ribavirin treatment significantly reduced viral titers and RNA levels, demonstrating antiviral sensitivity. Together, these findings provide new insights into BAGV biology, host adaptability, and potential zoonotic risk, and support the development of antiviral strategies and improved epidemiological surveillance within a One Health framework.

## INTRODUCTION

Bagaza virus (BAGV), first detected in 1966 in *Culex* mosquitoes in the Central African Republic(1), is an emerging flavivirus that has been regularly detected in wild birds between 2010 and 2023. Outbreaks have been reported in Spain (2010, 2019,2021)(2, 3), and in Portugal (2021, 2023)(4), frequently associated with disease and mortality, particularly in red-legged partridges (*Alectoris rufa,* 23-30% mortality)(5, 6), but also affecting other galliform and non-galliform wild bird species(7). Experimentally infected grey partridge (*Perdix perdix*) showed 40% mortality while sparrows were unaffected(8). Additional detections outside Europe, include the Himalayan monal (*Lophophorus impejanus*) in South Africa and mosquitoes in several African countries(9) and India(10). The increasing recurrence of detections over short periods in southern Europe, together with the geographic spread and expanding host range of BAGV, has raised concerns about its potential transmission to other wildlife species, domestic animals, and humans.

BAGV belongs to the genus *Orthoflavivirus*, which includes important human and animal pathogens such a dengue (DENV), West Nile (WNV), and Usutu (USUV). These single-stranded positive-sense RNA arboviruses are mainly transmitted by *Culex* or *Aedes* mosquitos(12). BAGV genome contains a single long open reading frame flanked by 5’-and 3’-untranslated regions (UTRs) and encodes a polyprotein that is post-translationally cleaved into three structural proteins, capsid (C), premembrane/membrane (prM/M), and envelope (E), and seven non-structural proteins, NS1, NS2A, NS2B, NS3, NS4A, NS4B, and NS5 proteins. Although this genomic architecture is highly conserved across flaviviruses, even subtle amino acid changes, particularly in E, NS2A, NS3, and NS5, can strongly influence host specificity, pathogenicity, and transmission dynamics. Among them, several avian-associated viruses, particularly WNV and USUV, have demonstrated the capacity to cross species barriers, occasionally causing neurological disease in humans and other mammals(13-15).

BAGV host range, replication dynamics, and adaptive capacity remain poorly understood, as most studies have focused on comparative genetics and outbreak epidemiology, with limited *in vitro* experimental evidence. BAGV mainly infects birds, but serological evidence in a human encephalitis case suggests potential zoonotic relevance(16). Although the BAGV 2010 strain was not infectious to adult mice in previous experimental trials(6), these findings highlight the adaptive potential of BAGV and the need to clarify its replication and host range within a *One Health* framework.

In this study, we report the successful isolation of infectious BAGV from avian tissues, followed by its adaptation to cell culture through serial passaging. We further characterized the replication kinetics across multiple host cell lines, identified adaptive mutations arising during culture, and described the cytoplasmic localization of replication intermediates. The sensitivity of this BAGV isolate to the broad-spectrum antiviral drug ribavirin was also tested in different cell lines, making available reference assays and tools to the identification of specific drugs against this putative zoonotic threat. Together, these results provide new insights into the replicative adaptability, evolutionary dynamics, and potential zoonotic capacity of BAGV, a virus that remains poorly characterized.

## MATERIALS AND METHODS

### Cell lines and culture conditions

Human (A549, HEK-293T, Huh7.5, LN229), mammalian (BHK-21, Vero), avian (CCL141, DF1) and mosquito (Aag2-AF5) cell lines were used as previously described(17). Briefly, mammalian and avian cells were cultured in DMEM (Sigma-Aldrich, St. Louis, USA) supplemented with 5–10% FBS (Sigma-Aldrich), 1% penicillin–streptomycin (Sigma-Aldrich) and L-glutamine (Gibco, UK). Aag2-AF5 cells were maintained in Leibovitz’s L-15 medium with glutamine (Gibco, USA) supplemented with 10% FBS (Gibco, USA), 1% non-essential amino acids (Gibco, USA), 1% penicillin–streptomycin (Sigma-Aldrich) and 10% tryptose phosphate broth (Gibco, USA). Cells were incubated at 37°C with 5% CO₂, except mosquito cells (28°C, no CO₂).

### Virus isolation

Tissue samples (encephalon, encephalon plus cerebellum, and growing feather) from a red-legged partridge found dead during the 2021 outbreak in Spain(3, 4) were used for virus isolation. Approximately 30 mg of tissue was homogenized in PBS using a mortar or rotor–stator homogenizer, centrifuged (900xg, 10 min, 4°C) and filtered (0.22 µm). BHK-21 and Vero cells were infected with 10, 50 or 100 µL of homogenate and blindly passaged every 3–7 days.

For RNA extraction, 30 mg of tissue was processed using the lysis buffer from GeneJET RNA Extraction Kit (Thermo Fisher Scientific) in the presence of 286 mM β-mercaptoethanol or 40 mM DTT. The tissues were homogenized in this buffer, using glass beads, followed by proteinase K treatment, and purification according to the manufacturer’s instructions. Purified RNA (10 µL) was transfected into BHK-21 or Vero cells using Lipofectamine 3000 (Thermo Fisher Scientific) in Opti-MEM as previously described(17, 18). Supernatants were collected at 48 h post-transfection and subjected to three successive blind passages in Vero cells at 7-day intervals, monitoring cytopathic effects. Viral presence was confirmed by conventional PCR, with cytopathic effects appearing progressively earlier upon passaging, and the highest viral titer obtained for BAGV passage 10 (BAGVp10) at 48 h post-infection (p. i.).

### Viral infection, replication kinetics and titration

Human, monkey, avian, and mosquito cells were infected at a multiplicity of infection (m.o.i). of 1 for replication kinetics and 3 for immunofluorescence experiments, as previously described(17, 18). Briefly, virus was adsorbed for 1 hour on cells cultured in DMEM with 1% FBS, and maintained in DMEM (1% FBS) until collection. Supernatants were harvested at 24, 48, 72, 96, and 120 h post-infection (p.i.) for titration.

Viral titers were determined by TCID₅₀ assays, as previously described(19). Briefly, 1×10⁴ Vero cells were seeded on 96-well plates in DMEM supplemented with 5% FBS (Sigma-Aldrich, St. Louis, USA). The next day, 10-fold serial dilutions of each viral sample were added. After 5 days, viral titers were calculated based on wells showing cytopathic effect using the Reed and Muench method(20).

### RT-PCR, genomic sequencing and analysis

RNA was extracted from supernatants of BAGV infected cells (48 h p.i.) using the GeneJET RNA Extraction Kit (Thermo Fisher Scientific) and reverse-transcribed into cDNA with random hexamers and Superscript III (Invitrogen). Viral fragments (∼3 kb) were amplified using Q5 high-fidelity DNA polymerase (New England Biolabs) and specific primers (Table 1) and annotated to the reference genome. The sequence of BAGV p10 was obtained by Sanger sequencing (STABVIDA, Portugal), while the sequences of BAGV p0 and BAGV WT were obtained by NGS sequencing (Catarina Fontoura-Gonçalves et al., unpublished). Sequence assembly and curation were performed with SeqMan (DNASTAR, Madison, WI, USA).

**Table 1.**
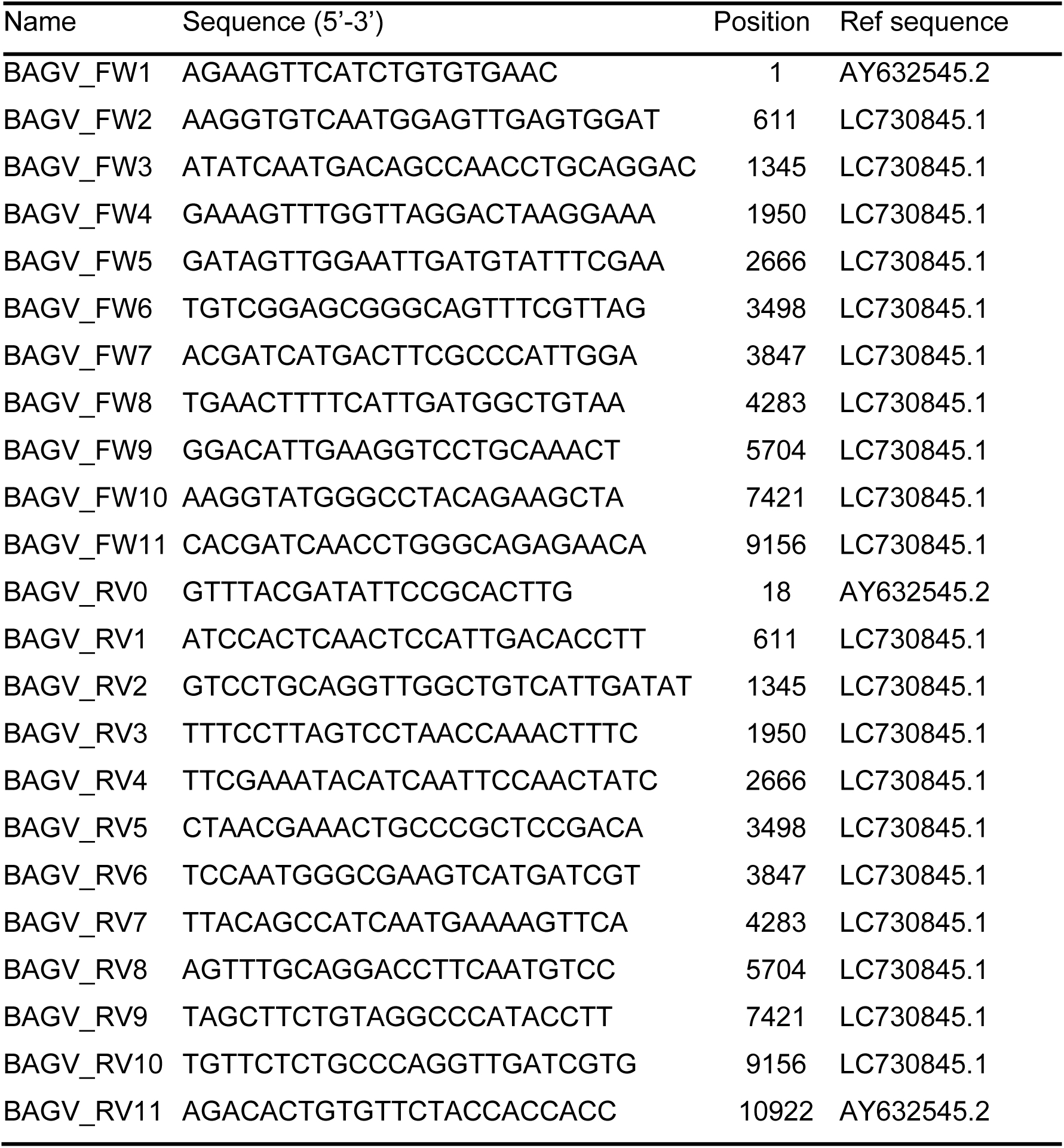
Oligonucleotides used in this study.

Genomes were compared to sequences available at GenBank, including those collected in previous Spanish outbreaks. FASTA sequences were aligned with Clustal Omega(21). PHYLIP-formatted alignments were used to construct neighbor-joining phylogenetic trees on the T-REX server(22), and visualized with ITOL(23). Selected 5`′` - and 3`′` - UTR regions were analyzed for secondary structure using MXfold2(24) and compared to WNV sequences available(GenBank M12294.2)(25).

### BAGV protein structure prediction and comparison

The amino acid sequences of the prM, E, NS1, NS2A, and NS3 proteins in BAGVp0 and BAGVp10 samples, predicted from genome sequencing, were used for in silico tertiary structure prediction using AlphaFold(26). Only structures with iPTM ≥0.8 were analyzed further. E, NS1, and NS3 proteins from both isolates were aligned with each other and also with WNV (E: PDB 2i69; NS1: PDB 4o6d; NS3 protease: PDB 8co8) and ZIKV (NS3 helicase: PDB 5jps) structures using UCSF ChimeraX to assess structural similarities(27).

### Immunofluorescence

To detect BAGV dsRNA intracellular, 1×10^5^ Huh-7.5 cells were seeded in 24 well culture plates. Immunocytochemistry was performed as previously described(28). Briefly, cells were infected at an m.o.i. of 3 and fixed at 48 h p.i. Then, dsRNA was detected using mouse anti-dsRNA (J2, Nordic Mubio) followed by Alexa 488-conjugated secondary antibody. Images were acquired as previously described(17, 28).

### Ribavirin treatment

Huh7.5, A549, and Vero cells (1×10⁵ per well) were infected at an m.o.i. of 1. After virus adsorption, cells were treated with ribavirin (1–200 µM) for 24 h. Supernatants were collected and viral titers determined as described above.

### Statistics

Data are expressed as mean ± SD. Comparisons between groups were analyzed by two-way ANOVA with Dunnett’s correction. Significance is indicated as ns (not significant), * (p<0.05), ** (p<0.01), *** (p<0.001).

## RESULTS

### Isolation and propagation of BAGV

To recover infectious BAGV from the tissues, we followed two different approaches in parallel: directly infecting cells with a sample rescued from homogenized tissue in PBS or transfection of cell cultures with RNA extracted from the same tissues. Those samples were obtained from three tissue types: growing feather, a mix of encephalon and cerebellum, and encephalon alone (Figure 1A).

**Figure 1.**
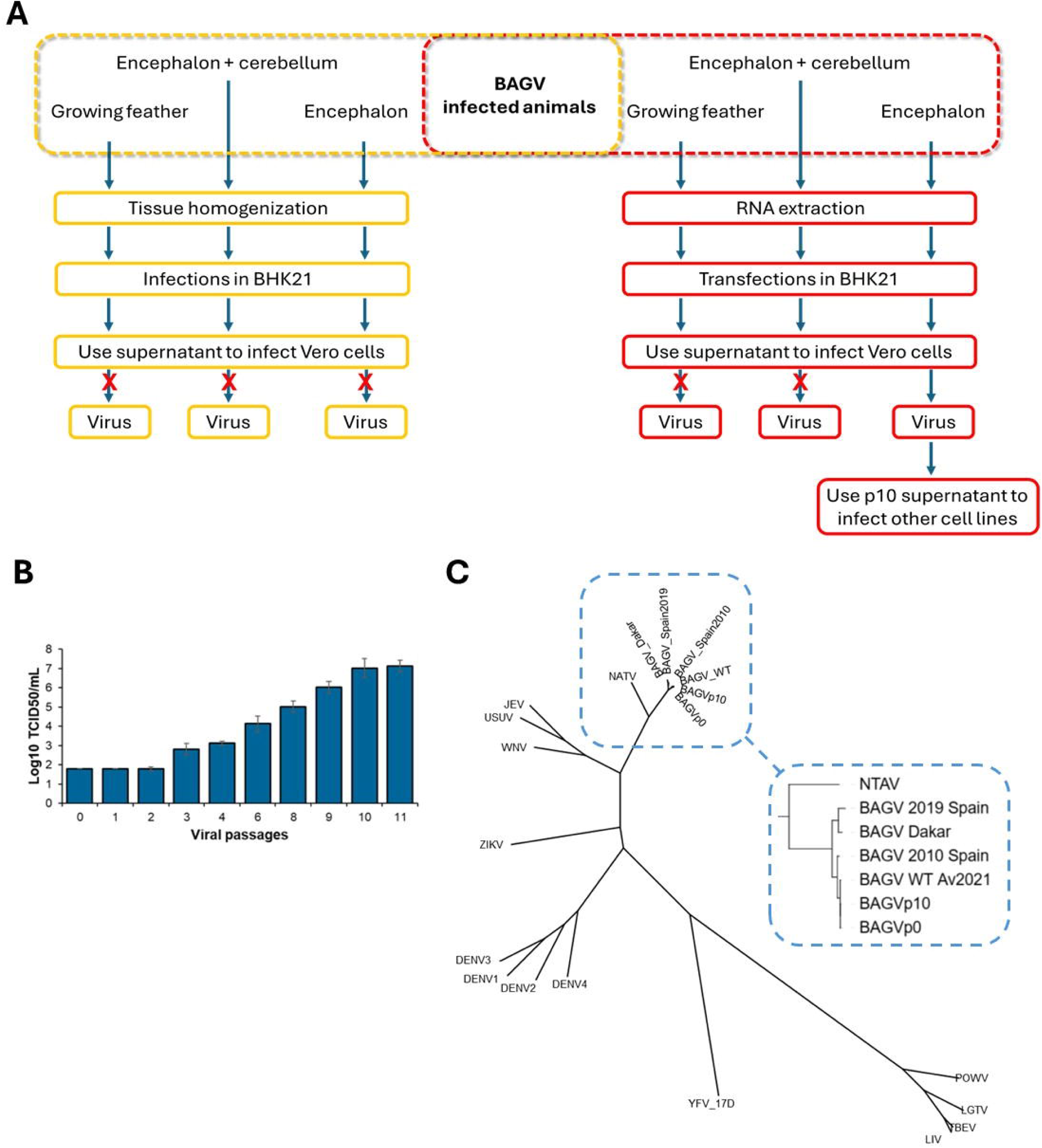
Isolation, adaptation, and phylogenetic analyses. **(A)** Schematic representation of the procedure used to obtain infectious viral particles in cell culture. Two different strategies were followed with BAGV-infected animal samples. The left part of the diagram (yellow) shows the strategy of direct infection of BHK-21 cells from homogenates of different tissues. The right side of the diagram (red) shows the strategy for total RNA extraction from the same infected tissues and subsequent transfection of this RNA into BHK-21 cells. In both cases, supernatants from BHK-21 cells were used to infect the Vero cells. Ten passages were performed in these cells, and the possible cytopathic effect (CPE) caused by the viral infection was analysed in each passage. Cytopathic damage was only observed in cells infected with material from the brain that followed the strategy of transfection with previously extracted RNA. The supernatant from passage 10 was expanded and saved for further experiments. **(B)** Viral titre in serial passages in Vero cells. The viral titer value (as Log_10_ TCID50/ml) obtained from the supernatant of serial passages in Vero cells is shown. The initial sample (0) corresponded to the supernatant of BHK-21 cells transfected with encephalon RNA, as described in Figure 1A. As can be seen from the graph, the first three passages were performed blindly as no cytopathic effect compatible with viral infection was observed, and a value of less than 1 × 10^2^ TCID50/ml was given. The passages were stopped when a plateau was reached between passes 10 and 11. **(C)** Phylogenetic analysis of isolates BAGV_WT, BAGVp0, and BAGVp10. The complete genome sequences of the isolates described in this study were compared with those of BAGV_Spain2010 (KR108244.1), BAGV_Spain2019 (PP236854.1), BAGV_Dakar (GCA_000883735.1), NATV (GCA_000897715.1), JEV (GCA_000862145.1), USUV (GCA_000854945.1), WNV (GCA_000861085.1), ZIKV (GCA_000882815.1), DENV(1-4) (GCA_000862125.1; GCA_000871845.1; GCA_000866625.1; GCA_000865065.1), YFV_17D (GCA_031123685.1), LIV (GCA_000863165.1), TBEV (GCA_000863125.1), LGTV (GCA_000860805.1), and POWV (GCA_000860485.1). The sequences were aligned using Clustal Omega, the alignment was converted to PHYLIP format, and the bootstrap analysis was performed on the T-REX web server (https://bio.tools/t-rex). The tree was rendered on the ITOL server (https://itol.embl.de). A clearer representation of the relationships between the BAGV isolates is shown in the enlarged box.

Direct inoculation of tissue homogenates onto BHK-21 or Vero cells did not result in detectable cytopathic effect (CPE), even when several blind passages of the cellular supernatants were carried out, regardless of the tissue and cell line used. In contrast, transfection of purified RNA from the encephalon into BHK-21 cells produced infectious progeny that could be then amplified in Vero cells. The supernatants from BHK-21 cells successfully infected Vero cells, where a clear CPE appeared after serial passaging, confirming active viral replication (Figure 1B). Neither RNA from growing feathers nor from the encephalon and cerebellum mix rescued productive infection. Three serial blind passages of Vero cells supernatants were required until the viral titer became detectable (Figure 1B). To successfully sequence the viral genome in early passages, which contained very low infectivity, the supernatants from passages 0 to 3 in Vero cells were pooled, and total RNA was extracted as described in the Materials and Methods section. This sample was named BAGVp0. The complete genome sequence of the virus recovered from passages 0 to 3 was obtained (BAGV_p0), along with the complete genome sequence of the virus extracted from the infected tissue (BAGV_WT), and the identity of BAGV was confirmed.

### Adaptation of BAGV to Vero cells

To evaluate the ability of BAGV to adapt to cell culture, the virus recovered from encephalon RNA was subjected to serial passaging in Vero cells. Viral titers progressively increased during passaging, reaching a plateau at passages 10-11, with values of approximately 1×10⁷ TCID₅₀/mL (Figure 1B). This confirms that BAGV undergoes efficient adaptation to Vero cells during the *in vitro* propagation.

To investigate the genetic changes associated with adaptation to the Vero cell line, we compared the consensus sequences of the original virus isolated from the feather (BAGV_WT), virus isolated from an early pooled passage (BAGVp0), and the virus after ten passages in Vero cells (BAGVp10). Phylogenetic analyses (Figure 1C) confirmed that these three isolates were closely related to the previously described BAGV_Spain2010 (2). We identified 17 synonymous and five non-synonymous mutations in the coding region, together with three additional changes located in the 3’-UTR (Figure 2A; Table 2). All the silent mutations were transitions (11 C→U and 6 A→G mutations), whereas non-synonymous mutations were three transitions and two transversions (Table 2). The 5’-UTR and coding regions corresponding to the proteins C and NS2B remained unchanged (Figure 2A).

**Figure 2.**
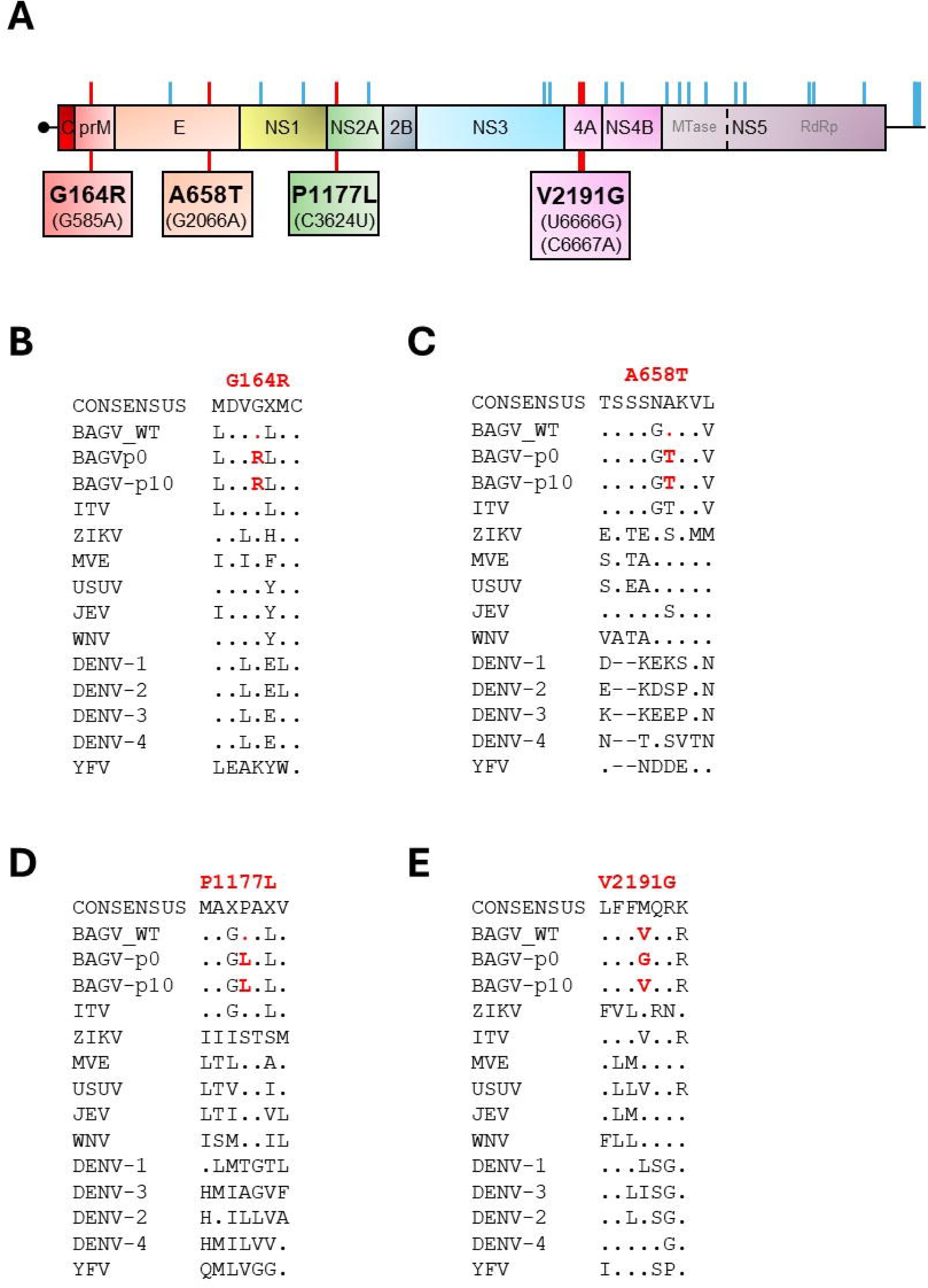
Localization of mutations acquired after 10 passages in Vero cells. **(A)** The BAGV genome map is shown with the untranslated areas at the 5‘ and 3’ ends and the coding region with each of the viral proteins: C, capsid; prM, premembrane; E, envelope; and NS1 to NS5, non-structural proteins 1 to 5. The schematic representation of the BAGV genomic structure also indicates the positions of the synonymous (blue) and non-synonymous (red) mutations described in Table 2, as well as the corresponding amino acid changes. **(B-D)** Amino acid sequence alignments comparing the BAGV isolates described in this work (BAGV_WT, BAGV_p0 and BAGV_p10) with the reference sequences of ITV, ZIKV, MVE, USUV, JEV, WNV, DENV-1, DENV-2, DENV-2, DENV-4, and YFV viruses for each mutated position described in this work (B: G164R; C: A658T; D: P1177L; E: V2191G). The consensus sequence is shown at the top of the figure. Dots in the alignment indicate identities with the consensus sequence. “*” indicates identity, while “:” and “.” indicate different degrees of similarity.

**Table 2.** Synonymous and non-synonymous mutations.

| Nucleotide position<sup>a</sup> | BAGV |  |  | Codon change | Amino acid change | Genome region |
| --- | --- | --- | --- | --- | --- | --- |
|  | WT | BAGVp0 | BAGVp10 |  |  |  |
| 584 | G | A | A | GGA→AGA | **G164R** | prM |
| 1603 | U | C | C | AAU→AAC |  | **E** |
| 2066 | G | A | A | GCA→ACA | **A658T** |  |
| 2788 | C | U | U | GAC→GAU |  | NS1 |
| 3226 | U | U | C | CAU→CAC |  |  |
| 3624 | C | U | U | CCA→CUA | **P1177L** | **NS2A** |
| 4030 | U | C | U | UUU→UUC |  |  |
| 6170 | U | Y | U | UUG→CUG |  | NS3 |
| 6245 | A | G | G | AAA→AAG |  |  |
| 6666 | U | G | U | GUC→GGA | **V2191G** | **NS4A** |
| 6667 | C | A | C |  |  |  |
| 6931 | G | G | A | AAG→AAA |  | NS4B |
| 7138 | U | C | U | GGU→GGC |  |  |
| 7693 | C | U | U | AUC→AUU |  | NS5-MTase |
| 7840 | A | R | A | GCA→GCG |  |  |
| 7951 | U | U | C | GUU→GUC |  |  |
| 8176 | A | G | G | UUA→UUG |  |  |
| 8536 | C | U | U | ACC→ACU |  | NS5-RdRp |
| 8683 | G | G | A | GUG→GUA |  |  |
| 9430 | C | U | U | AUC→AUU |  |  |
| 9436 | A | G | G | GUA→GUG |  |  |
| 10103 | U | C | C | UUG→CUG |  |  |
| 10743 | U | C | C | n.a.<sup>b</sup> |  | 3'-UUR |
| 10769 | A | G | G | n.a. |  |  |
| 10785 | U | C | C | n.a. |  |  |

<sup>a</sup>Position referred to the BAGV-WT isolate (GenBank reference number: PX907777).
<sup>b</sup>n.a. not applicable. GenBank reference number BAGVp0: PX907778 y BAGVp10: PX907776.

Substitutions accumulated predominantly within the non-structural protein-coding region and the 3’-UTR (22 out of 25 mutations), suggesting that early adaptation involved selective pressure acting mainly outside the structural gene block. Non-synonymous changes became fixed in BAGVp10 in prM (G164R), E (A658T), NS2A (P1177L), and NS4A (V2191G), indicating that mutations at these amino acid positions were likely associated with increased replicative fitness in Vero cells. In contrast, no amino acid substitutions were found in C, NS1, NS2B, NS3, NS4B, or NS5. Notably, some differences emerged when comparing the early and late passages. BAGVp0 contained a transient transversion in NS4A (V2191G) that was absent from the BAGV_WT sequence and reverted during subsequent passages, as it was absent in the BAGVp10 sequence (Table 2).

Comparison with published BAGV genomes revealed that the G164R change in prM affects a residue that is highly conserved across *Orthoflavivirus* genus, with BAGVp0 and BAGVp10 acquiring a charged amino acid, similar to that observed in yellow fever virus (YFV) (Figure 2B). The remaining substitutions in E, NS2A, and NS4A mapped to less conserved positions and corresponded to amino acids found with variable frequencies across other flaviviruses (Figures 2C–2E). No alterations were detected in the conserved domains implicated in enzymatic activity. Altogether, these data show that BAGV undergoes genetic diversification during cell culture adaptation and that only a small, specific subset of mutations becomes fixed as the virus acquires an enhanced replicative capacity in Vero cells.

### Non-synonymous mutation structure predictions

As no high-resolution structures or 3D models of BAGV proteins are available, the structural impact of the identified mutations was assessed by comparison with evolutionary related (homologous) flavivirus proteins. The absence of BAGV-specific structural data limits residue-level mapping but does not preclude inferences based on conserved domains and motifs that are well-characterized across the *Orthoflavivirus* genus. The structure prediction of the prM, E, NS2, and NS4A proteins was obtained using the AlphaFold program, but only the E protein showed high confidence metrics (PTM) (equal to or greater than 0.8) compared to the crystal structures of WNV. In addition, the results showed a high overlap of the compared structures (Figure 3A, blue WNV, green BAGV). The WNV structure used for this comparison (PDB ID: 2i69) is glycosylated at position N443 (Figure 3A, red), and this residue was conserved in the BAGV isolate (Figure 3B). Residue 658 is in loop 3 of domain III (D-III) of the envelope protein, near the glycosylation site, in a highly conserved region (Figure 3B). The A658T mutation alters the H-bonding pattern of this loop. A Thr residue at this position can create H-bonds with the backbone oxygen at position S655 and with the side chain of T653, likely stabilizing the loop (Figure 3A).

**Figure 3.**
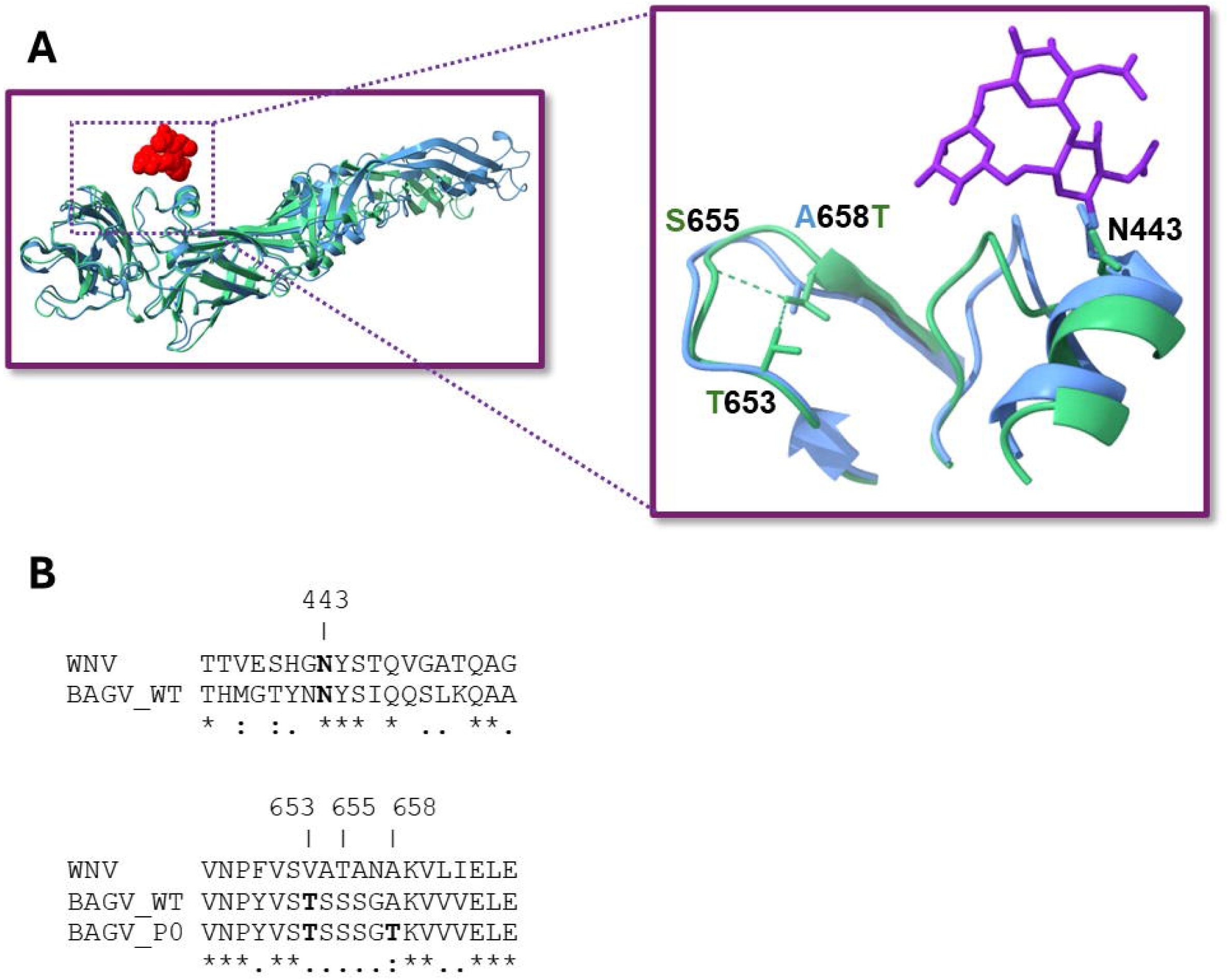
Protein structure comparison. **(A)** Alignment of the structure of the E protein of BAGVp0 (PTM = 0.81) with that of the E protein of WNV (PDB ID: 2i69) using AlphaFold. On the right is a zoom of loop3, where the A658T mutation is located, and the hydrogen bridges that are modified because of this change. (**B)** Alignment of the amino acids in the E protein region, where the N-glycosylation site (N443) is located (top), and the A658T mutation at the bottom. The symbols at the bottom of the alignments (* : .) mean no change, very conserved, and conserved, respectively.

### Untranslated region mutations

Differences in the 5`′` and 3`′` UTRs were examined by comparison with the WNV genome, the most closely related orthoflavivirus with available UTR structure`–`function data. In the 5`′` UTR, the SLA and SLB regions showed moderate to high identity and similarity values of 49.3%/74% and 69.2%/84.6%, respectively, and exhibited notable differences in predicted secondary structure. In particular, the start codon is in a paired region in WNV but within a loop in the BAGV stem–loop structure (Figures 4B and 4C).

**Figure 4.**
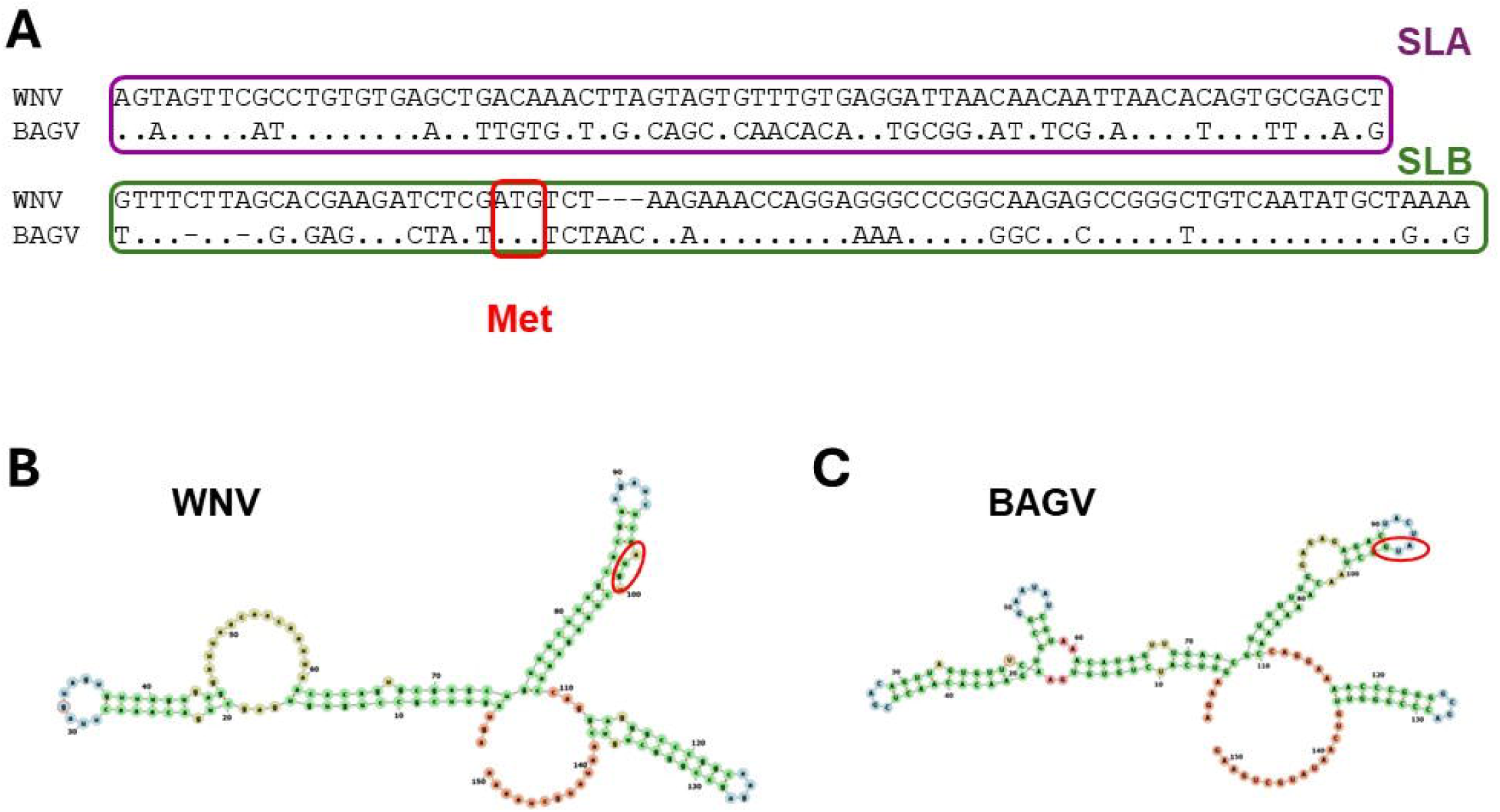
Sequence alignment of the 5’UTR. **(A)** Sequence alignment of the 5’-UTR region of BAGV_WT was compared to the corresponding WNV sequence. The sequences corresponding to the SLA and SLB regions are marked in magenta and green, respectively. Dots indicate the same nucleotide as the upper sequence. Gaps introduced to maintain the alignment are indicated by dashes. The start codon (AUG) is indicated in red. (**B)** and (**C**) The sequence of the 5’UTR region shown in part A was used for secondary structure prediction using the MXFold2 software(24). Secondary structure predictions of the 5’UTR regions of WNV (**B**) and BAGV_WT (**C**) are shown in the figure.

In contrast, alignment of the 3′UTR genomic sequences revealed a high degree of conservation between BAGV and WNV, with no major differences in the SL-I to SL-IV and 3′SL regions (Figure 5A). However, a large deletion affecting the 5′DB-RCS2 region and the WNV PK3 pseudoknot caused substantial changes in the predicted secondary structure, while the terminal sHP and 3′SL regions remained highly conserved (Figures 5B and 5C).

**Figure 5.**
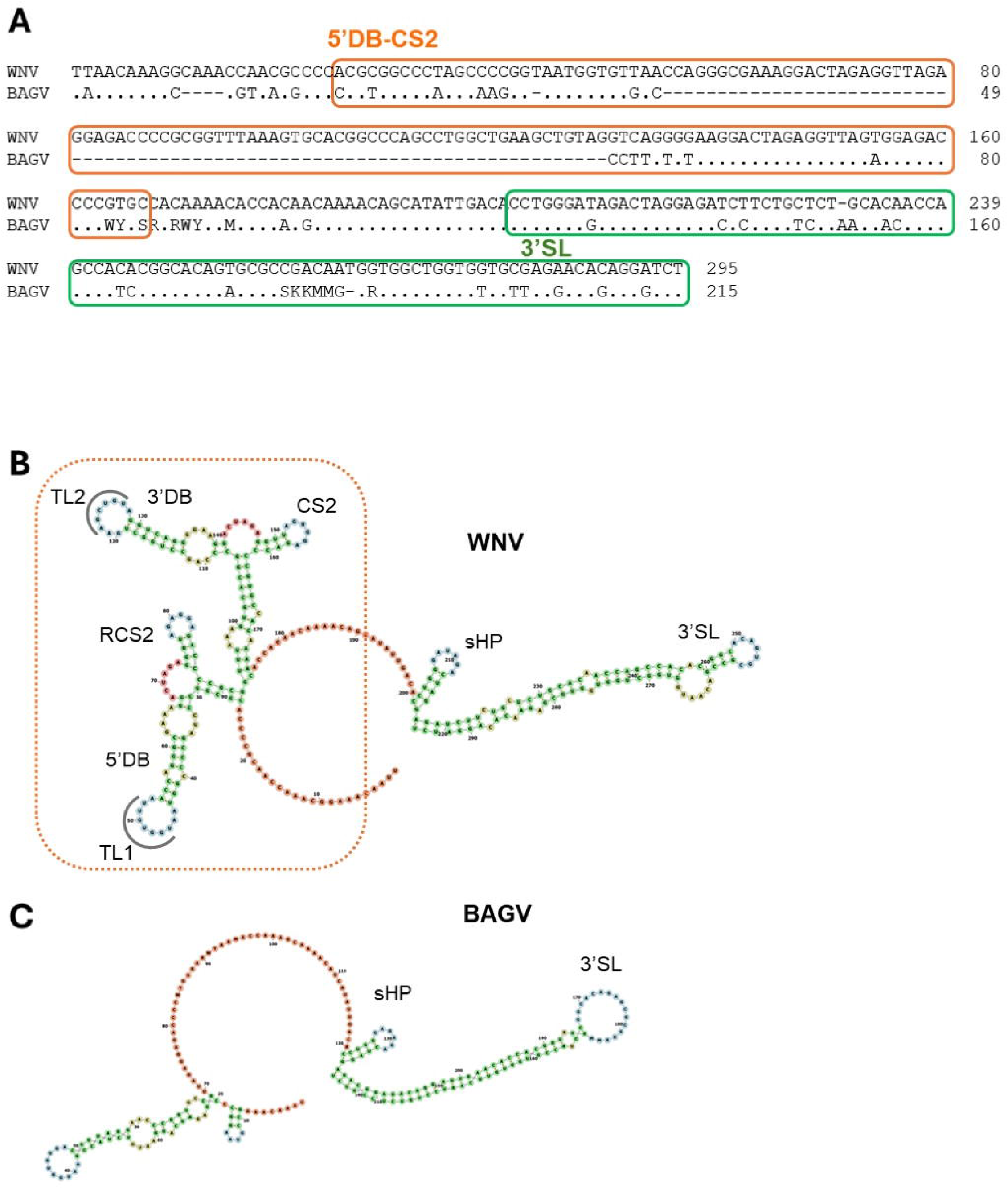
Sequence alignment of the 3’UTR. **(A)** Sequence alignment of the 5’-UTR region of BAGV_WT compared to the corresponding WNV sequence. Sequences corresponding to the 5’DB-CS2 and 3’SL regions are marked orange and green, respectively. Dots indicate the same nucleotide as the upper sequence. Gaps introduced to maintain the alignment are indicated by dashes. (**B)** and (**C)**. The sequence of the 3’UTR region shown in part A was used for secondary structure prediction using the MXFold2 software(24). Secondary structure predictions of the 3’UTR regions of WNV (**B**) and BAGV_WT (**C**) are shown in the figure. The regions TL1, 5’DB, RCS2, TL2, 3’DB, CS2, sHP, and 3’SL are indicated as described in(25, 34). The missing sequence in BAGV is indicated by a dotted orange box.

### Replication kinetics in different cell lines

Following adaptation of BAGV to Vero cells, replication kinetics were evaluated in mammalian, avian, and mosquito cell lines (Figure 6). BAGV replicated efficiently in monkey (Vero), mosquito (Aag2-AF5), and duck (CCL-141) cells, as well as in human HEK-293T (kidney) and Huh7.5 (liver) cells. Vero cells were among the most permissive, with titers rapidly increasing to ∼7.6 log at 48 h p.i. and remaining high at least until 120 h p.i. Aag2-AF5 mosquito cells showed highly efficient replication, reaching peak titers of ∼7.7 log at 96 h. Duck CCL-141 cells displayed moderate replication during the first 48 h, followed by an increase to ∼6.1 log at 72 h, after which further time points could not be analysed. In human HEK-293T and Huh7.5 cells, high early titers (∼7.0 and ∼6.8 log, respectively) were followed by a pronounced decline at later time points.

**Figure 6.**
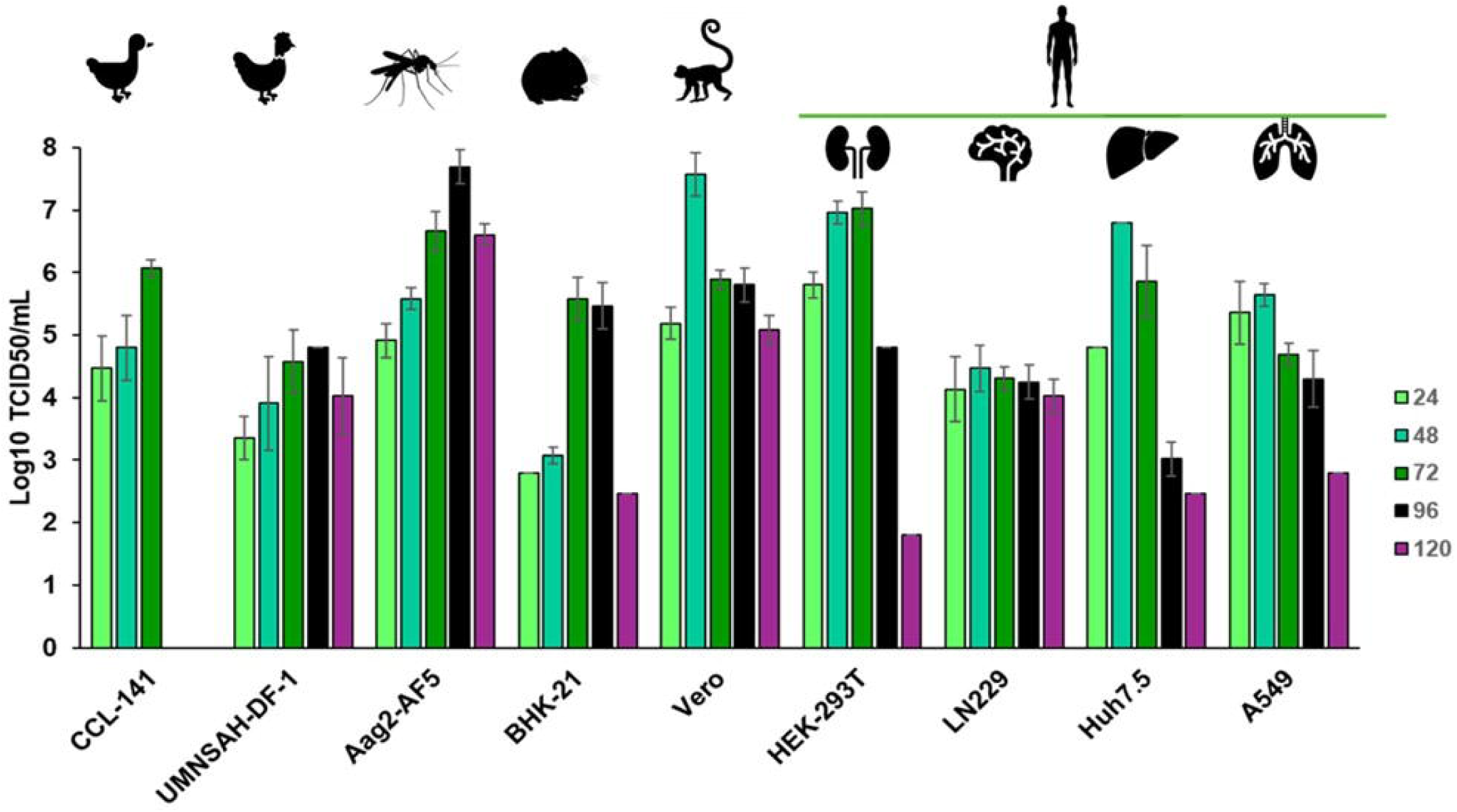
Viral titer in cell lines of different origin. The viral titer obtained after infection of CCL-141 (duck), UMNSAH-DF-1 (chicken), Aag2-AF5 (mosquito), BHK-21 (hamster), Vero (monkey), HEK-293T (kidney, human), LN229 (glioblastoma, human), Huh7.5 (liver, human) and A549 (lung, human) cell lines are shown. Cells were infected with BAGV propagated in Vero cells up to passage 10 (Figure 2). Culture supernatants were collected at 24, 48, 72, 96, and 120 h post-infection.. These supernatants were used for virus titration, as described in the Materials and Methods section. The results shown correspond to the mean of two biological replicates, each in triplicate, and their corresponding standard deviations.

In contrast, viral titres were lower in human A549 and LN229 cells, hamster BHK-21 cells, and chicken UMNSAH-DF-1 fibroblasts. Of those, viral replication kinetics in LN229 cells were relatively stable throughout the experiment, with titers of ∼4.0–4.5 log TCID_50_/ml. Overall, viral titers in different cell lines generally reached a maximum value between 48 and 72 h p.i., except in mosquito cells, which peaked at 96 h p.i.

### Subcellular localization of BAGV replication complexes

To assess the subcellular localization of BAGV replication, dsRNA distribution was analysed by confocal microscopy in BAGVp10 infected and uninfected cells. Infected cells showed a punctate accumulation of dsRNA restricted to the cytoplasm, with a prominent perinuclear distribution, consistent with flavivirus replication complex formation (Figure 7A). No dsRNA signal was detected in uninfected controls, confirming that BAGV replication is exclusively cytoplasmic and follows a typical flaviviral replication pattern.

**Figure 7.**
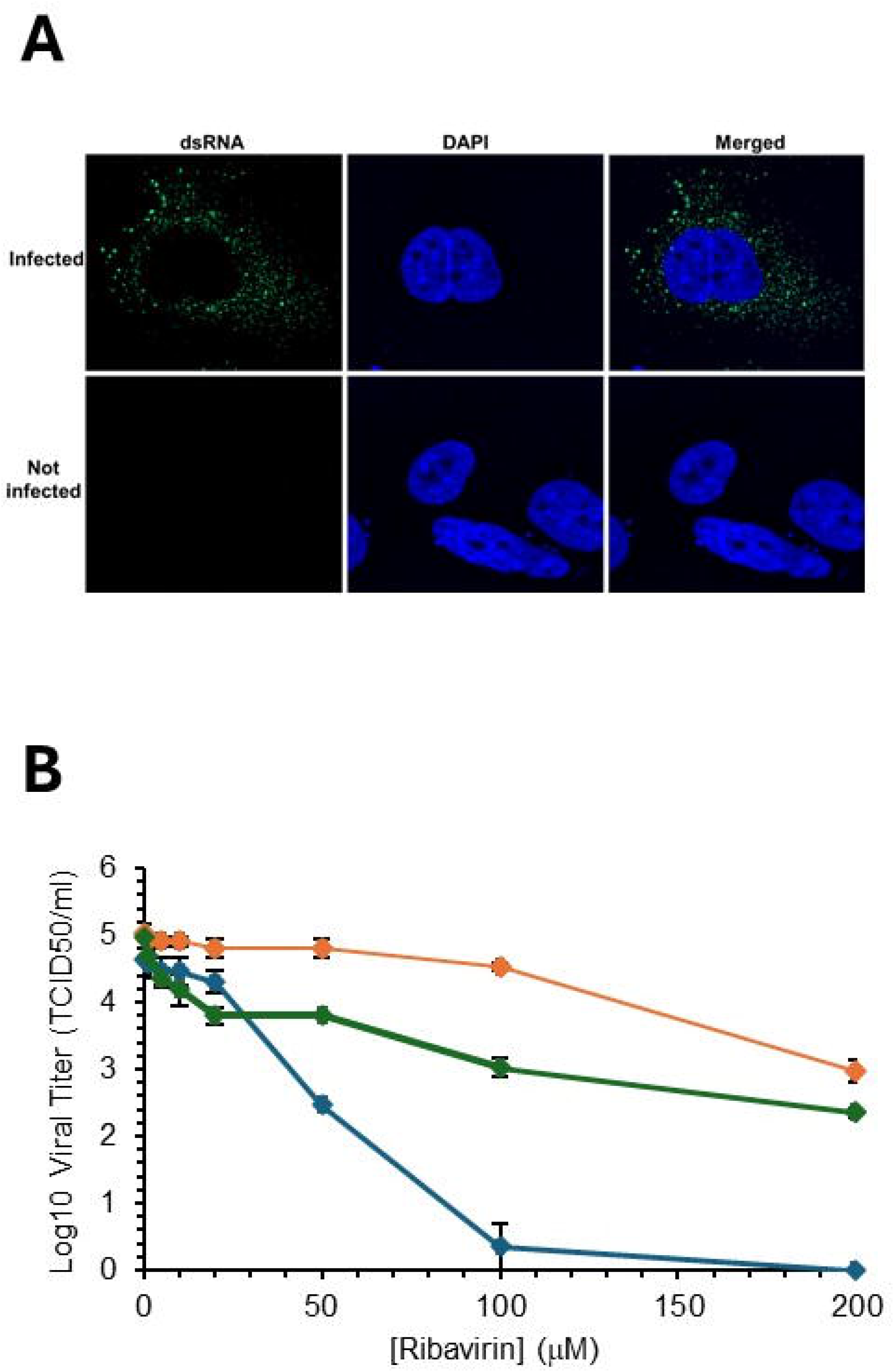
Replication complexes and treatment with ribavirin. **A**. Images from immunocytochemistry experiments showing localization of double-stranded RNA (green) and nucleus (blue) in infected (top) and uninfected (bottom) cells are shown. **B**. Effect of ribavirin treatment of cells of different origin infected with BAGVp10. The viral titer (expressed as TCID50/ml) of Huh7.5 (orange), A549 (green), and Vero (blue) cells treated with ribavirin at the indicated concentrations at each point on the graph is shown.

### Response of BAGV to ribavirin treatment

To assess BAGV sensitivity to ribavirin, Huh7.5, A549, and Vero cells were infected with BAGVp10 and treated with increasing ribavirin concentrations (0–200 µM). Viral titers decreased significantly in all cell types in a dose-dependent manner (Figure 7B), with reductions of ∼2 logs in Huh7.5 cells, 2.6 logs in A549 cells, and ≥4.6 logs in Vero cells, below the detection limit.

## Discussion

BAGV is an emerging orthoflavivirus in southern Europe responsible for recurrent outbreaks in wild birds. However, the lack of experimental systems and baseline biological data has severely limited risk assessment, surveillance, and preparedness efforts. Given its zoonotic potential and relevance for animal and wildlife health, a better understanding of BAGV biology is needed. We therefore isolated BAGV by transfecting total RNA into Vero cells and evaluated its replication in human, animal, and mosquito cell lines. As a result, the adapted virus (BAGVp10) accumulated synonymous and non-synonymous mutations and replicated in multiple host cells with variable efficiencies and ribavirin sensitivity.

Virus isolation was achieved exclusively through RNA transfection, as direct inoculation of tissue homogenates failed (Figure 1A), likely due to low levels of infectious particles or sample degradation, as previously reported for BAGV and other flaviviruses(6). This approach enabled the recovery of infectious virus from encephalon samples and the establishment of a productive culture system (Figure 1B). These observations align with prior reports of BAGV isolation in Spain and Portugal (2, 4, 6), although none of those studies explored the differential success of inoculation versus RNA-based rescue.

After isolation, BAGV rapidly adapted to Vero cells, reaching stable titers of ∼1×10⁷ TCID₅₀/mL after ten passages. Full-genome sequence analysis identified 17 synonymous and 5 non-synonymous mutations, along with three 3′-UTR changes, indicating diversification at both coding and non-coding levels during cell culture adaptation (Table 2; Figure 2). Several of these mutations have been reported in Iberian isolates, supporting their recurrence and functional relevance(3). Notably, the prM G164R substitution replaces the glycine present in European genotype 2 isolates from the 2010 and 2021 outbreaks(2, 3) with the arginine found in genotype 1 African strains(29), suggesting strong functional constraint and repeated selection across genotypes. Additional non-synonymous changes in E, NS2A, and NS4A occurred at less conserved positions within the *Orthoflavivirus* genus but have been observed in other flaviviruses (Figures 3C–3E), potentially influencing viral fitness by affecting protein stability, membrane association, or polyprotein processing. Moreover, although often considered neutral, synonymous substitutions can modify RNA secondary structure, codon usage, and replication efficiency(30-33). Consistently, the predicted secondary structure of the 3’-UTR revealed differences between BAGV_WT, BAGVp0, and BAGVp10, suggesting that RNA-level constraints may also contribute to fitness gains during passaging(25, 34).

BAGV replication kinetics varied markedly among cell lines, revealing differences in host susceptibility and cell line immune response. Robust replication was observed in mosquito Aag2-AF5 and mammalian Vero, HEK-293T, and Huh7.5 cells, with distinct peak kinetics between insect and mammalian systems likely reflecting differences in antiviral responses, metabolism, and temperature-dependent replication(11, 35-38). Replication in avian cells was generally lower, with higher titers in duck-derived CCL-141 than in chicken DF-1 cells, consistent with host-specific differences reported for other avian flaviviruses(13, 15, 17, 35). Among human cell lines, LN229 glioblastoma cells showed limited susceptibility, although this does not exclude neurotropism, as this line is poorly permissive to several neurotropic flaviviruses(17). The detection of BAGV in avian encephalons(4) and a reported human meningoencephalitis case(16) highlight the need to further assess BAGV infection in CNS-associated cell models in future studies.

The ecological context of BAGV supports its potential for cross-species transmission. In Europe, BAGV has been detected in several bird species, primarily red-legged partridges, ring-necked pheasants (*Phasianus colchicus*), wood pigeons (*Columba palumbus*), and more recently Eurasian magpies (*Pica pica*), among others, indicating an expanding host range(2, 4, 39). Vector competence studies show that *Culex tritaeniorhynchus* and *Aedes aegypti* can efficiently support BAGV replication, including high viral loads in saliva and transovarial transmission, supporting its spillover potential to other vertebrates, including humans(10). Phylogenetic analyses further indicate that European BAGV isolates form a distinct cluster from West African lineages, consistent with multiple introductions followed by local persistence and ecological adaptation(3, 7, 39, 40).

Finally, we demonstrate that ribavirin exhibits antiviral activity against BAGV, although the degree of inhibition varies depending on the cell type used. This variability is consistent with the known cell line-dependent action of ribavirin against other flaviviruses and highlights the importance of selecting appropriate in vitro systems for antiviral assessment(41-43). The BAGVp10 isolate described here provides a robust model for future antiviral screenings. Given that ribavirin is a broad-spectrum inhibitor(44), additional nucleoside analogs, such as favipiravir, sofosbuvir, and galidesivir, which have been previously shown to inhibit related flaviviruses, should be evaluated to identify more potent candidates.

We established a BAGV isolate that replicates efficiently in diverse cell lines and reaches high viral titers. This adaptation is driven by the selection of specific mutations, including highly conserved amino acid changes across BAGV genotypes, underscoring their importance for viral fitness. Identifying these conserved positions provides insight into adaptation mechanisms and supports optimization of virus isolation and propagation from field samples. Moreover, the efficient replication of BAGV in multiple cell types establishes this isolate as a valuable tool for structure–function studies, antiviral screening, and the development of attenuated strains. Overall, this study advances our understanding of BAGV molecular adaptation, host range, and drug susceptibility, providing a critical experimental foundation for assessing its potential impact on animal and human health within a *One Health* framework.

## Acknowledgements

The authors gratefully acknowledge Dr. Kevin Maringer for generously providing Aag2-AF5 mosquito cells. Work in the laboratory of AM is supported by AEI funding (grant numbers PID2019-106068GB-I00 and PID2022-137974OB-I00) and JCCM funding (grant number SBPLY/21/180501/000076). LAG and AA are the recipients of grant SBPLY/24/180225/000240 from Junta de Comunidades de Castilla-La Mancha (JCCM). JQ is supported by FCT Invited Research Chair in Hunting and Biodiversity. CFG is supported by an FCT PhD grant (reference 2022.12139.BD). We also acknowledge the UCLM grant numbers 2022-GRIN-34150 and 2025-GRIN-38349,, the BAGA-PT (2022.09263.PTDC; https://doi.org/10.54499/2022.09263.PTDC) project, from the Fundação para Ciencia e Tecnologia (FCT), Portugal, and the technical assistance of J.J. Lorenzo. This research was conducted within the framework of a research stay funded by the University of Castilla-La Mancha (Spain), through the *Call for Research Stays at Universities and Research Centres Abroad for the year 2025 (Second term) (Resolution of 6 November 2024) BDNS (Identif.): 795356. [2024/9041]*.

